# Diversity of RNA viruses in declining Mediterranean forests

**DOI:** 10.1101/2023.10.28.564537

**Authors:** Sergio Diez-Hermano, Pedro Luis Pérez-Alonso, Wilson Acosta Morell, Jonatan Niño-Sanchez, Marcos de la Peña, Julio Javier Diez

## Abstract

Global change alters forestry habitats and facilitates the entry of new pathogens that don’t share a co-evolution history with the forest, leading them into a spiral of decline. As a result, relationships between forests’ organisms get disbalanced. Under this scenario, RNA viruses are of particular interest, as they participate in many of such relationships thanks to their ability to infect a wide range of hosts, even from different kingdoms of life. For this reason, the study of RNA viruses is essential to understand how viral flow across different hosts might occur, and to prevent possible outbreaks of diseases in the future. In this work the RNA virus diversity found in trees, arthropods and fungi from declining Mediterranean forests is described. To this extent, three habitats (*Quercus ilex, Castanea sativa* and *Pinus radiata*) were sampled and RNAseq was performed on tree tissues, arthropods and fungi. 146 viral sequences were detected by searching for matches to conserved motifs of the RNA-dependent RNA polymerase (RdRP) using Palmscan. Up to 15 viral families were identified, with Botourmiaviridae (28.7%) and Partitiviridae (9.6%) being the most abundant. In terms of genome type, ssRNA(+) viruses were the most represented (83.5%), followed by dsRNA (15%) and two ssRNA(-) representatives. Viruses belonging to families with cross-kingdom capabilities such as Hypoviridae (1), Mitoviridae (6) and Narnaviridae (5) were also found. Distribution of viruses across ecosystems was: *Q.ilex* (57.5%), *P.radiata* (26.7%) and *C.sativa* (15.8%). Interestingly, two RdRP sequences had no matches in available viral databases. This work constitutes a starting point to gain insight into virus evolution and diversity occurring in forests affected by decline, as well as searching for novel viruses that might be participating in unknown infectious pathways.

## 1 Introduction

In recent decades, it has been shown that all biotic constituents of forest ecosystems can carry and be affected by viruses. In addition to plant viruses, all plant-associated organisms (bacteria, protists, fungi, vertebrate and invertebrate animals) have their own viral communities (virome), and together they form the metavirome or virosphere of the forest system. Currently, 64 virus species are known to affect 19 forest plant species in temperate and urban forests (Rumbou et al., 2021). Based on genome type (single-stranded or ss, double-stranded or ds, positive-sense, or antisense), most are (+)ssRNA viruses belonging to the *Beta-, Seco-, Bromo-, Tombus-, Virga-* and *Mayoviridae* families. In comparison, only one (-)ssRNA family has been found so far, the *Emaraviridae*, consisting of four species. Three dsRNA viruses have also been described in pine and ash, as well as three other dsDNA viruses with retrotranscriptase capacity in *Betula* sp., *Castanea sativa* and *Fraxinus americana*. Among all the viral diversity described, the genera *Emaravirus* and *Badnavirus* are considered to be tree pathogens and, therefore, causative agents of the diseases with which they are commonly associated.

In contrast to trees’ virome, most viruses affecting forest fungi and oomycetes have a dsRNA genome and belong to the *Partiti-, Toti-, Curvula*- and *Reoviridae* families. Viruses with a (+)ssRNA genome are classified within the families *Hypo-, Endorna-, Botourmia-, Fusari-* and *Mitoviridae*, and some of their members have been found to be able to induce changes in the level of pathogenicity of forest pathogens of particular relevance, such as *Cryphonectria parasitica, Ophiostoma novo-ulmi, Gremmeniella abietina, Fusarium circinatum, Heterobasidion annosum* and *Hymenoscyphus fraxineus* (van Diepeningen, 2021). In addition, the first (-)ssRNA and ambivalent (combining + and - sense strands in their genome) viruses of fungal pathogens have recently been described in *Armillaria* sp. and *C. parasitica* (Forgia et al., 2021; Linnakoski et al., 2021). Finally, high viral diversity has also been found in soil pathogens such as *Rosellinia necatrix*, leading to the discovery of new *Hypoviridae* and candidates yet to be classified, such as *Fusagraviruses* and *Megatoviruses* (Arjona-López et al., 2018).

Current virus taxonomy is based on the presence of homologous genomic structures between viral families. In particular, plant viruses tend to cluster with fungal and arthropod viruses (Lefeuvre et al., 2019). This is indicative of the ability of some viruses to overcome barriers between biological kingdoms and infect a wide diversity of organisms (cross-kingdom viruses), which, in the context of forests, mainly involves trees, fungi and insects. Cross-kingdom viruses represent a considerable proportion of the fungal virome, to the extent that 50% of known fungal species carry plant viruses (Cao et al., 2022). Some members of the *Toti-, Partiti-, Endorna*- and *Chrysoviridae* families are plant viruses capable of replicating in meristematic cells and infecting fungi in vitro (Nerva et al., 2017). The first experimental evidence of a natural cross-kingdom infection was found in *Rhizoctonia solani*, which was able to acquire and transmit the Cucumber mosaic virus from its host plant (Andika et al., 2017). Fungal and plant viruses are able to act synergistically and facilitate transmission even between vegetatively incompatible fungal strains, using the plant as a transport pathway. This phenomenon has been described experimentally for the *Cryphonectria hypovirus-1* and the Tobacco mosaic virus, and active expression of the p29 gene of the former and the movement proteins (MPs) of the latter were necessary for it to take place (Bian et al., 2020). MPs are unique to plant viruses and are necessary for viral particles to move from cell to cell. However, fungal viruses, which do not usually have an extracellular phase, lack them. This display of functional cooperation gives an idea of the potential of viruses to infect unexpected organisms, regardless of the evolutionary distance that separates them.

Insects are also susceptible to cross-kingdom viral infections. Despite being the most numerous group of vectors transmitting plant viruses, few studies have been devoted to unravelling the virome of insects affecting forest species. Once inside the insect, plant viruses can develop forms of non-persistent transmission (in external structures of the insect) or circular transmission (using internal compartments as reservoirs) (Whitfield et al., 2015). In some cases, circular transmission may involve replication of the plant virus within the cells of the insect host. In the case of European forests, viruses have been found in *Lepidoptera* such as *Lymantria dispar, Thaumetopoea pityocampa* and *Leucoma salicis*, in *Coleoptera* such as *Ips typographus* and in *Hymenoptera* such as *Neodiprion sertifer* (Jakubowska et al., 2015; Paraskevopoulou et al., 2021). Fungal viruses capable of infecting insects, such as some DNA viruses of the *Genomoviridae* family, have also been detected (Liu et al., 2016).

Yet, despite having more information than ever about the viral component of forest ecosystems, studies on the virosphere of Mediterranean forests are still lacking, to the best of our knowledge. Mediterranean forests rank among the terrestrial ecosystems threatened by globalisation and climate change the most (Newbold et al., 2020). Weather predictions for the coming decades project drastic reductions in precipitation patterns and rising temperatures, exposing forests in the Mediterranean basin to long periods of drought and massive forest fires (Nunes et al., 2022). The high frequency of climatic adversities decreases the ability of Mediterranean tree species to cope with the high number of pests and diseases spread by globalisation, facilitating the invasion of new pathogens with which forests have not had the opportunity to co-evolve. There is evidence that most RNA viruses, many of which are known causative agents of emerging infectious diseases, are contained in forest ecosystems and are often released as a result of forest disturbance and expanding human populations (Wilcox and Ellis, 2006; Mackay and Arden, 2016), increasing the risk that unexpected pandemics might arise.

Therefore, given the capacity of RNA viruses to jump between organisms and even to alter essential characteristics such as the pathogenicity level of their host, getting a better understating of the viral flows that might occur in endangered ecosystems is of the essence. With this purpose, the metavirome of three Mediterranean forests affected by decline is described here, by means of high-throughput sequencing and for different kingdoms of life, including plants, fungi and arthropods.

## 2 Materials and methods

### 2.1 Sampling sites and procedure

Bark, wood and leaves’ samples were extracted from trees of *Castanea sativa* (chestnut), *Quercus ilex* (holm oak) and *Pinus radiata* (Monterrey pine) from Castile and Leon (Spain) forests (Table 1 and Figure 1). Each sampling site corresponded to a single tree species. Sampling was conducted for all four sites in June and July 2021. Material from sixteen trees was sampled and pooled per plot. After removing the external bark, one sample was taken per tree from the main trunk at the height of 50 cm over the collar, to a depth of 2-3 cm. Only xylem and the internal bark layer (phloem) were considered in the analysis. Living individuals of common arthropods (ants, grasshoppers, beetles) were also collected (8-10 specimens each). Samples were stored at 4°C for one day prior to processing and at -80°C afterwards.

**Figure 1.**
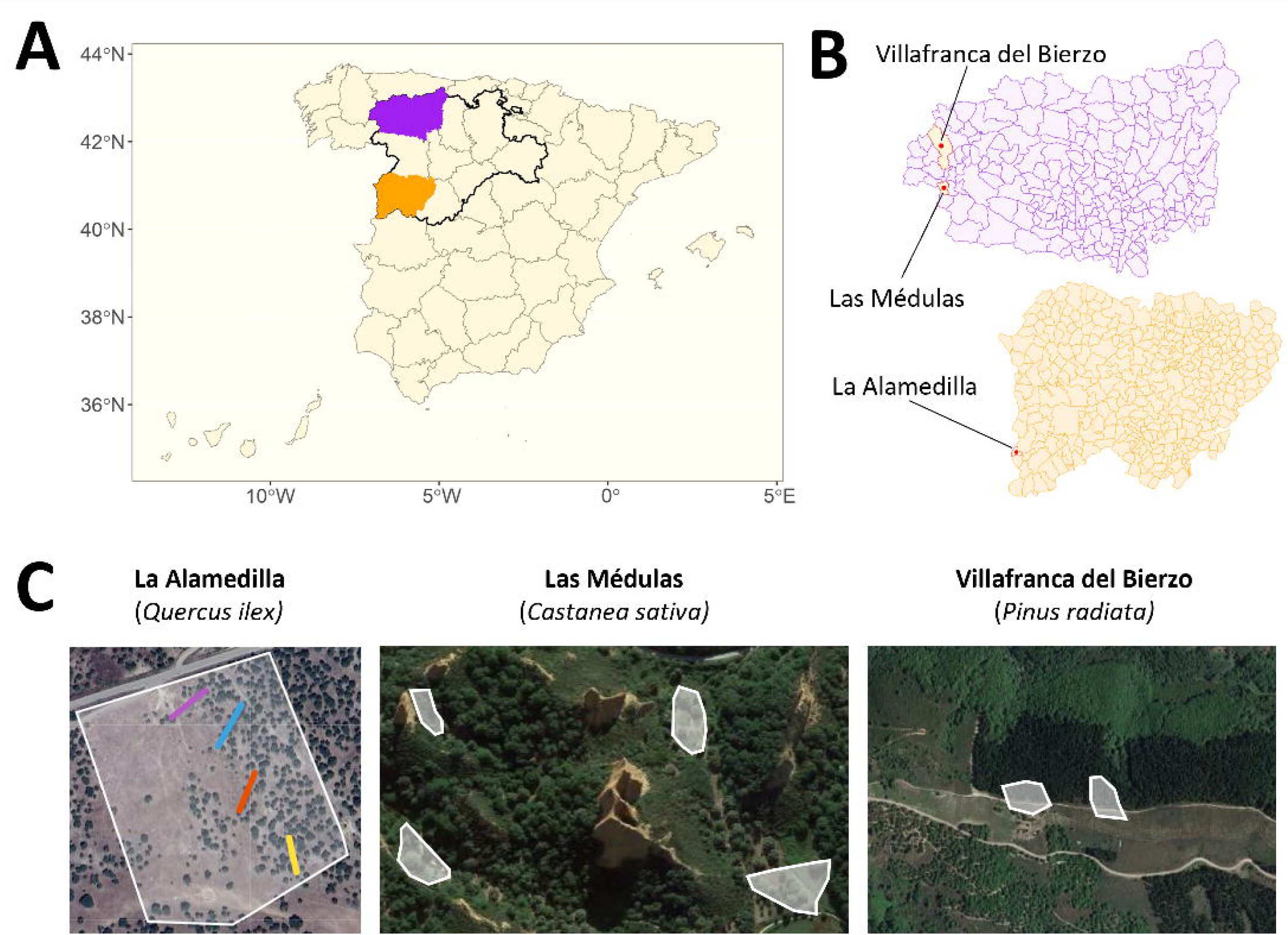
Map of sampling sites. **A** Map of Spain. Castile and Leon region is highlighted with a black and thick contour. Orange and purple areas correspond to Salamanca and Leon provinces, respectively. **B** Municipalities from where samples were taken (top: Leon, bottom: Salamanca). **C** Sampling areas (white polygons) overlayed on top of physical maps. Coloured lines in La Alamedilla correspond to four sampling transects.

**Table 1.**
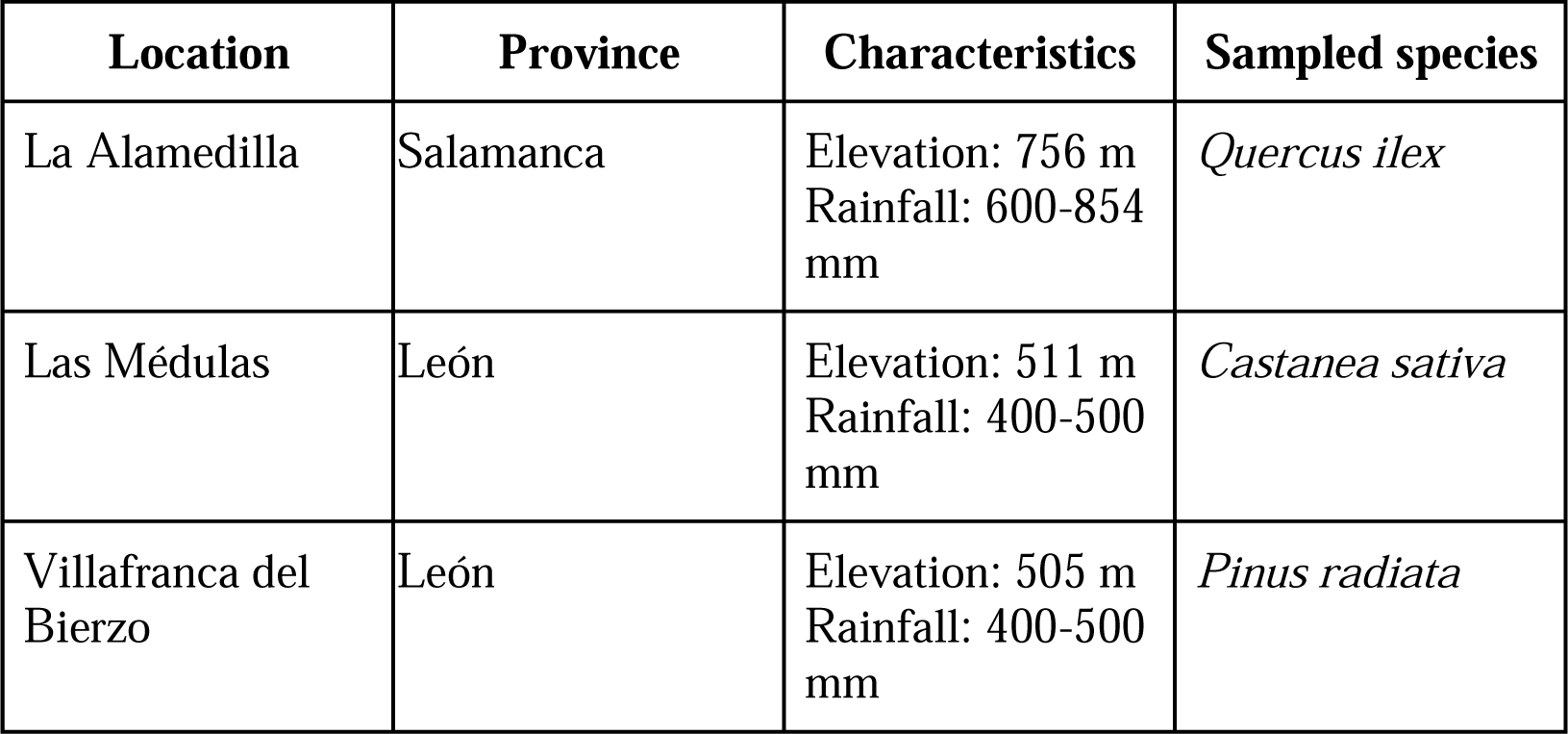
Description of sampling sites.

| Location | Province | Characteristics | Sampled species |
| --- | --- | --- | --- |
| La Alamedilla | Salamanca | Elevation: 756 m<br>Rainfall: 600-854 mm | *Quercus ilex* |
| Las Médulas | León | Elevation: 511 m<br>Rainfall: 400-500 mm | *Castanea sativa* |
| Villafranca del Bierzo | León | Elevation: 505 m<br>Rainfall: 400-500 mm | *Pinus radiata* |

### 2.2 Sample processing and sequencing

Fungi were cultured from the trees and arthropods’ samples. Bark, wood and leaves were cut into small pieces and disposed on Petri dishes with PDA+ampicilin (100 ug/ml) medium for 3 days at 25°C. Arthropods were washed in a Ringer solution and 50 ul of the solution were poured and spread also on Petri dishes with PDA+ampicilin medium for 3 days at 25°C. Visually different fungal colonies were picked, isolated and stored at -80°C.

Frozen samples of bark, wood, leaves and arthropods were pulverised and pooled per sampling site prior to RNA extraction. Isolated fungi were pooled likewise. Total RNA was purified from trees, arthropods and fungi using Spectrum Plant kit (Sigma Aldritch, MO, USA) following the manufacturer’s protocol. Quality control with Qubit yielded RIN values > 7 for all extractions. Samples were sent to Macrogen (South Korea) for sequencing (Illumina Miseq with TruSeq libraries and ribosomal RNA depletion, 150 bp paired-end reads).

### 2.3 Bioinformatic analysis

Raw reads were cleaned and trimmed using Cutadapt v.3.5 (-q 20) (Martin, 2011). Contigs were assembled with SPAdes v.3.15.5 (rnaviralSPAdes.py) (Prjibelski et al., 2020) and scanned for the presence of RNA-dependent RNA polymerase (RdRp) sequences using Palmscan v1.0 (Babaian & Edgar, 2022), a software that searches sequencing data for matches with the conserved motifs A, B and C (the “palmprint”) from the primary protein sequence of RdRp. RdRp sequences that matched any non-viral organism in nr NCBI database were discarded. Viral identities were assigned to the remaining RdRps by aligning the found sequences against PALMdb (Edgar et al., 2022) and RdRp-scan (Charon et al., 2022) databases using DIAMOND v2.1.8 (Buchfink et al., 2021). Taxonomy resolution was stablished according to different percent identity thresholds (species at >90%, genus at >70% and family at >30%). Multiple sequence alignment was performed with MAFFT (E-INS iterative refinement method) (Katoh et al., 2019) and used as input for phylogenetic tree building on IQTREE (default parameters) (Nguyen et al., 2015) with ultrafast bootstrap (Hoang et al., 2018) and automatic model selection (Kalyaanamoorthy, 2017).

Viral diversity analyses were performed and visualised in R environment 4.2.1 (R Core Team, 2022) using the packages *ggtree* (Yu, 2020), *ggmsa* (Zhou et al., 2022) and *seqinr* (Charif et al., 2023). Sequence logos were plotted with Skylign (Wheeler et al., 2014)

## 3 Results

### 3.1 Taxonomy and distribution of RdRps

A total of 146 RdRps were found among all analysed samples. Of these, 122 (83%) were assigned to existing viral groups with different levels of resolution, 22 (15%) had matches in current databases but lacked taxonomy assignment and 2 were absent from any database (Table 2 and Table 3). Most of the RdRps were found in the habitat of *Q. ilex* (57.5%), followed by *P. radiata* (26.7%) and *C. sativa* (15.8%) (Figure 2 and Table 4). Regarding sample type, half of the RdRps were found in tree samples (46.5%), followed by arthropods (21.2%), fungi from arthropods (16.4%) and fungi from trees (15.7%).

**Figure 2.**
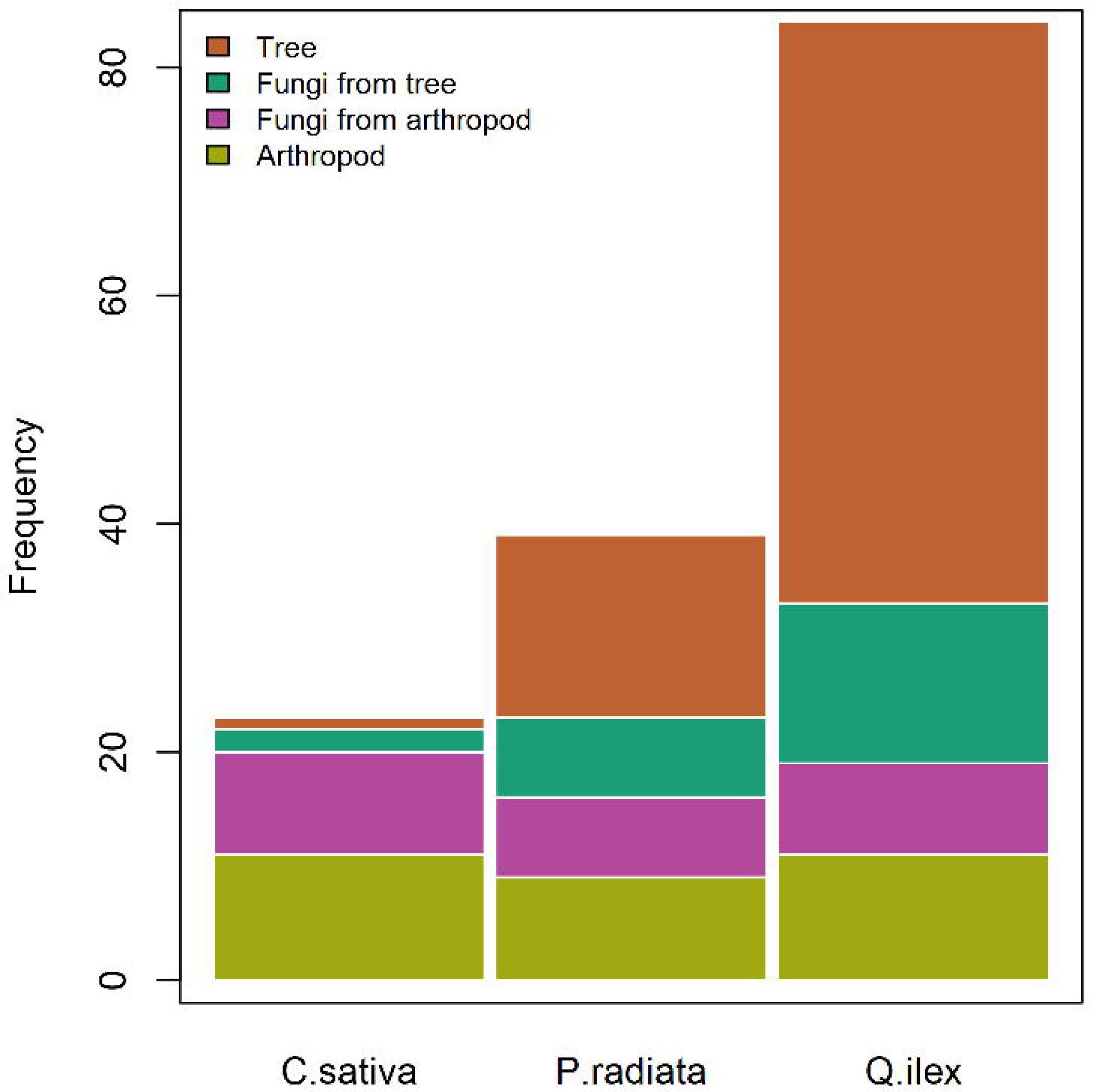
RdRp counts by habitat and sample type.

**Table 2.**
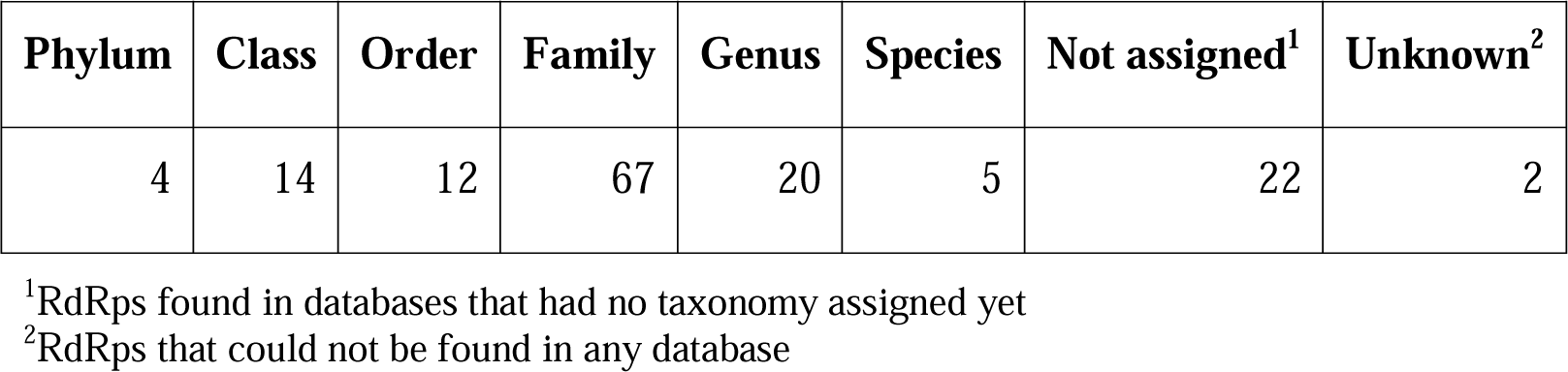
Taxonomic distribution of RdRps.

| Phylum | Class | Order | Family | Genus | Species | Not assigned<sup>1</sup> | Unknown<sup>2</sup> |
| --- | --- | --- | --- | --- | --- | --- | --- |
| 4 | 14 | 12 | 67 | 20 | 5 | 22 | 2 |

<sup>1</sup>RdRps found in databases that had no taxonomy assigned yet
<sup>2</sup>RdRps that could not be found in any database

**Table 3.**
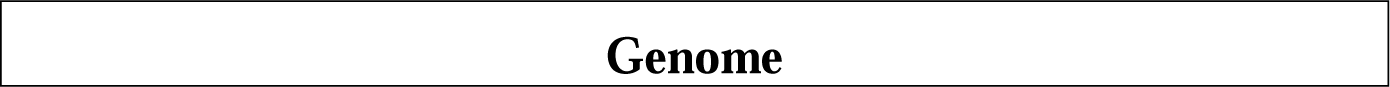
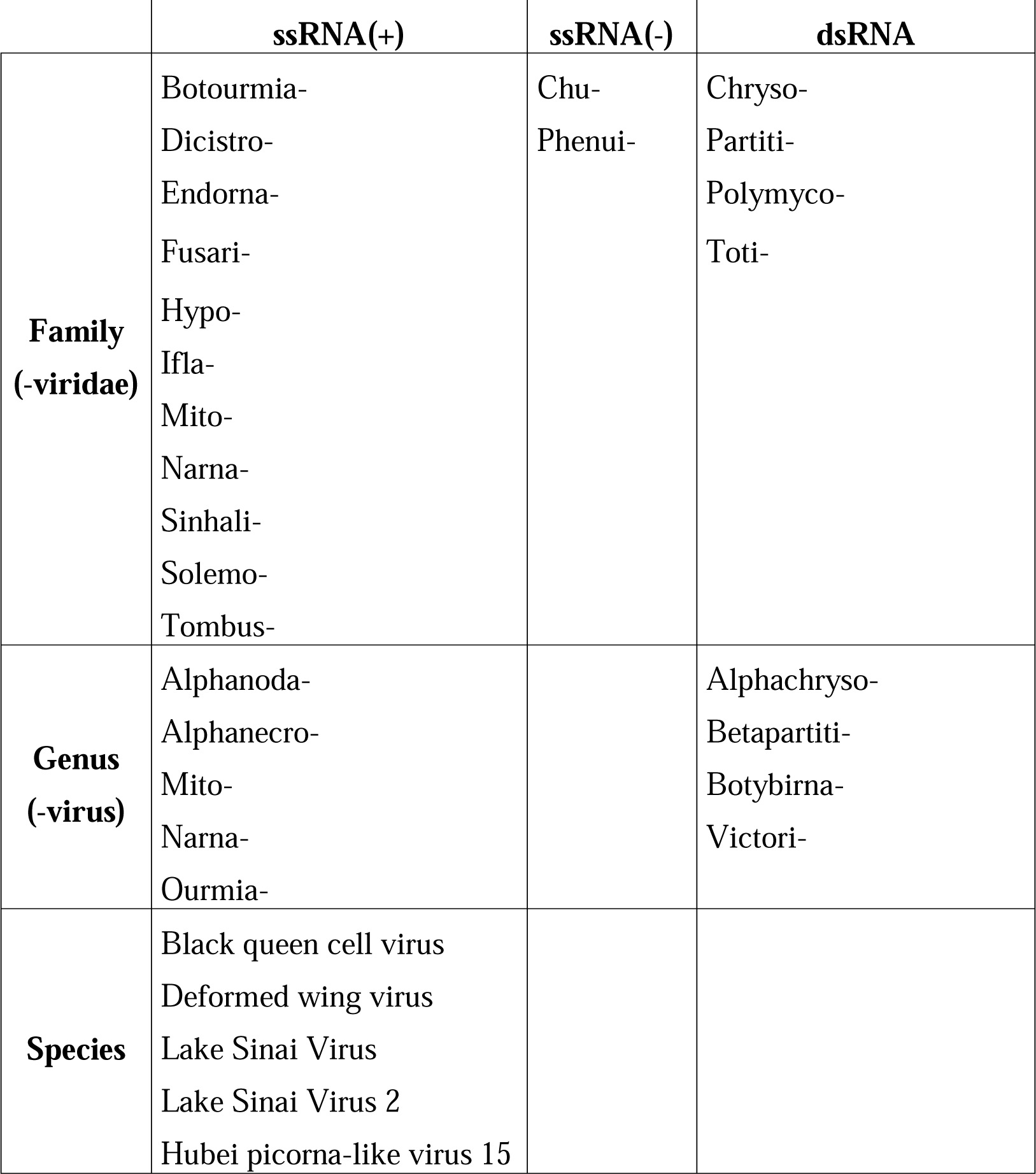
Taxonomic assignment of RdRps per genome type.

|  | ssRNA(+) | ssRNA(-) | dsRNA |
| --- | --- | --- | --- |
| **Family**<br>**(-viridae)** | Botourmia-<br>Dicistro-<br>Endorna-<br>Fusari-<br>Hypo-<br>Ifla-<br>Mito-<br>Narna-<br>Sinhali-<br>Solemo-<br>Tombus- | Chu-<br>Phenui- | Chryso-<br>Partiti-<br>Polymyco-<br>Toti- |
| **Genus**<br>**(-virus)** | Alphanoda-<br>Alphanecro-<br>Mito-<br>Narna-<br>Ourmia- |  | Alphachryso-<br>Betapartiti-<br>Botybirna-<br>Victori- |
| **Species** | Black queen cell virus<br>Deformed wing virus<br>Lake Sinai Virus<br>Lake Sinai Virus 2<br>Hubei picorna-like virus 15 |  |  |

**Table 4.** Count of RdRps per habitat and sample type.

|  | Tree | Fungi from tree | Arthropod | Fungi from arthropod |
| --- | --- | --- | --- | --- |
| **Chestnut** | 1 | 2 | 11 | 9 |
| **Pine** | 16 | 7 | 9 | 7 |
| **Holm oak** | 51 | 14 | 11 | 8 |

The most numerous viral families were Botourmiaviridae (28.7%), Partitiviridae (9.6%), Mitoviridae (4.1%) and Narnaviridae (4.1%) (Figure 3). In terms of genome type, ssRNA(+) viruses were the most represented (83.5%), followed by dsRNA (15%) and two ssRNA(-) representatives, Chuviridae and Phenuiviridae, found in arthropods from pine and chestnut habitats, respectively.

**Figure 3.**
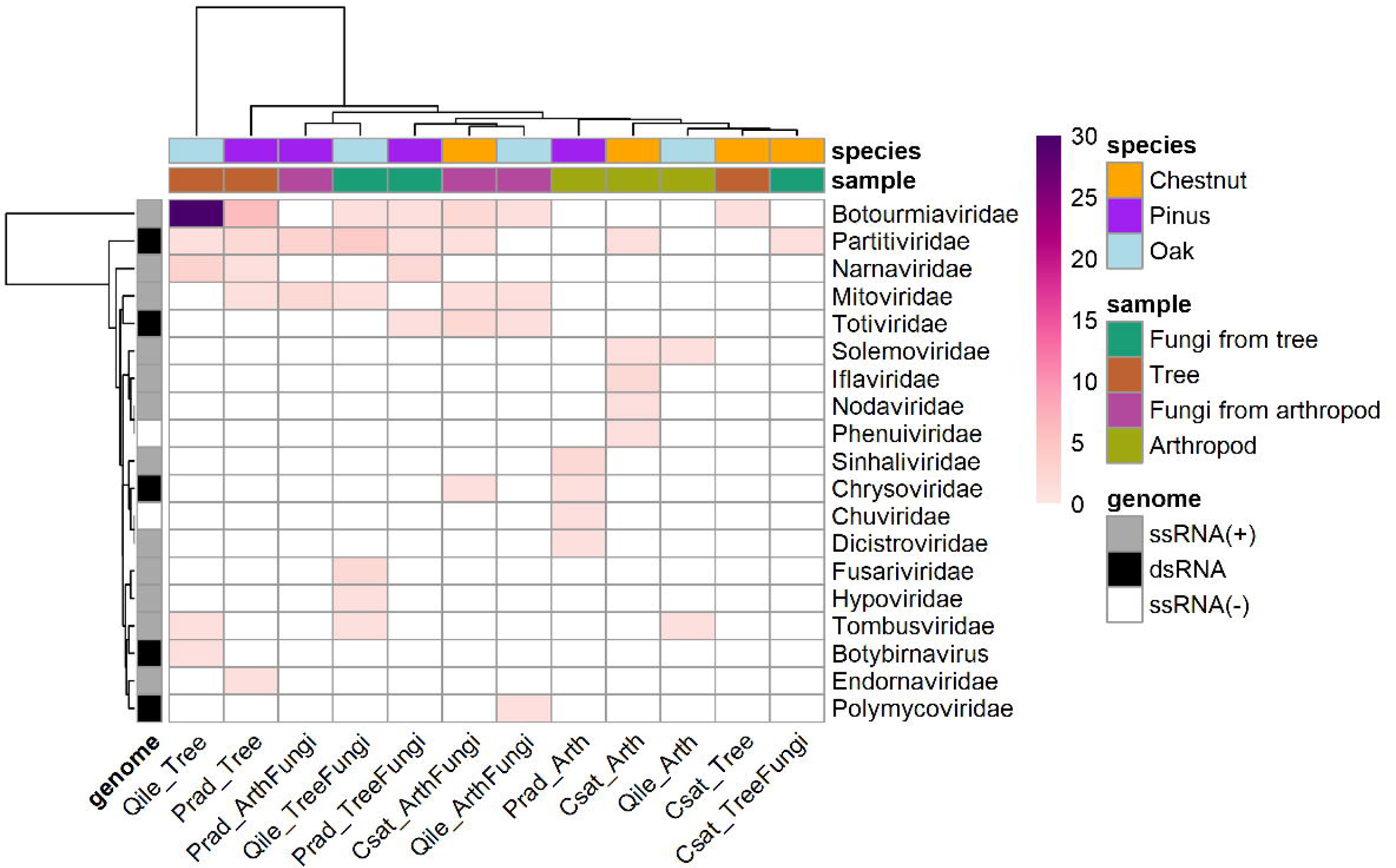
Distribution of viral families per habitat, sample type and genome type. Colour’s gradient from white to purple indicates abundance.

The distribution of RdRps revealed that sample types have different viral profiles (Figure 3). Some of the families were exclusive to fungi (Totiviridae, Fusariviridae, Hypoviridae and Polymycoviridae) or arthropods (only in chestnut habitat: Iflaviridae, Nodaviridae, Phenuiviridae; and only in pinus habitat: Sinhaliviridae, Chuviridae and Dicistroviridae). One family was found to be exclusive to holm oak’s habitat (Tombusviridae), regardless of sample type. Endornaviridae family was found only in pines whereas Botybirnavirus was exclusive to holm oaks.

### 3.2 Phylogeny of RdRps

MSA of the RdRps resulted in the dendrogram shown in Figure 4. RdRps from trees and arthropods’ samples clustered in two distinct groups, while RdRps from isolated fungi were interspersed, forming heterogeneous subgroups. No grouping by habitat was observed except for most RdRps from holm oak samples clustering together within the Lenarviricota supernode. All of the RdRps identified up to species levels belonged to arthropods’ samples: Lake sinai virus 1 and 2, Deformed wing virus, Black queen cell virus and Hubei picornalike virus 15, in the Kitrino- and Pisuviricota nodes. In nodes where two RdRp from trees’ fungi and arthropods’ fungi were paired, as is the case for some Betapartitivirus and Narnaviridae (bottom part of the tree), their sequences shared 100% identity. In all other cases the homology in the conserved RdRp motifs A, B and C was at least greater than 50%.

**Figure 4.**
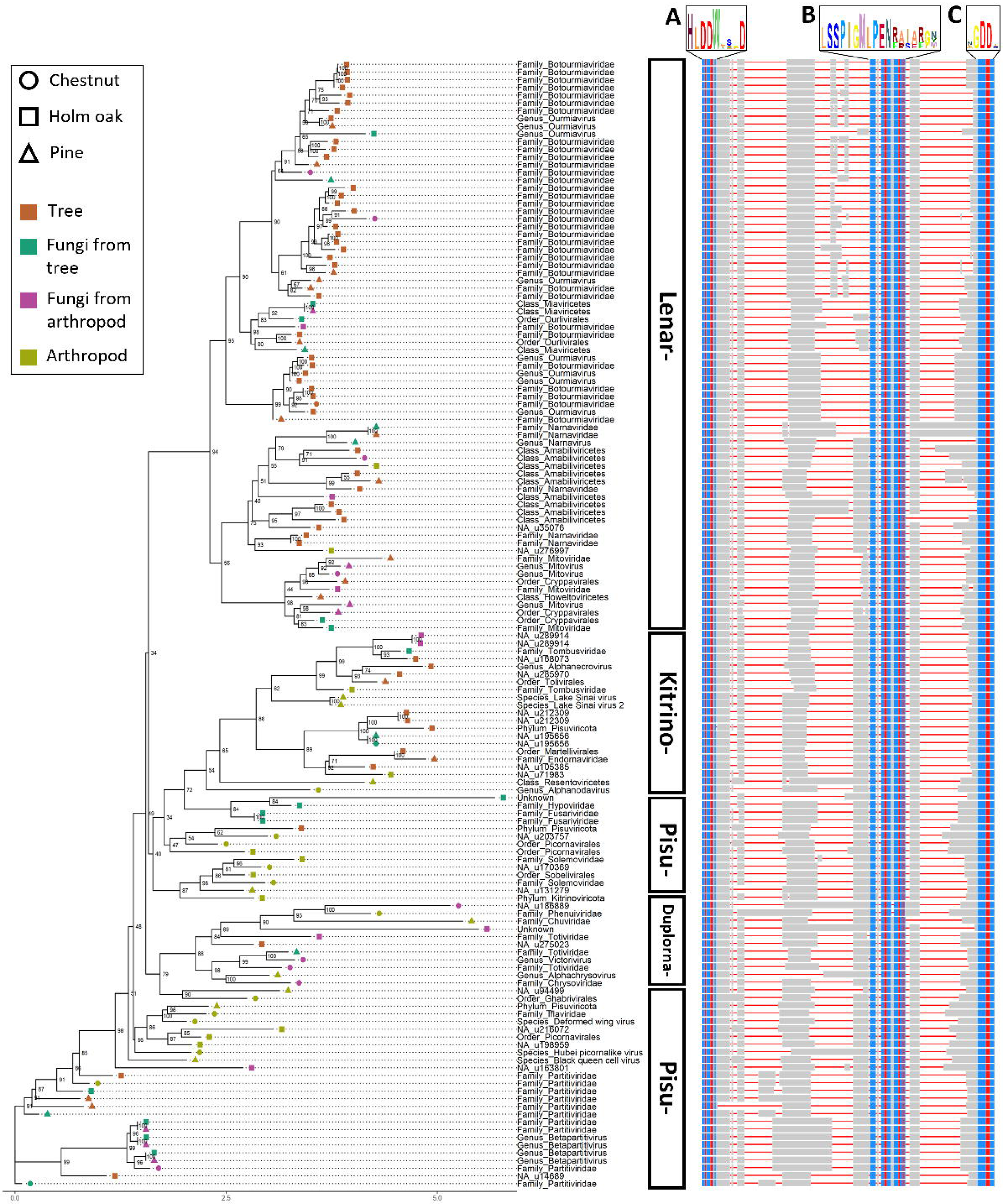
Phylogenetic tree of RdRps. Left: consensus tree with bootstrap confidence values. Each branch is labelled with the taxonomic level followed by the identity assigned to that particular RdRp. Vertical rectangles with black outline indicate viral phyla (-viricota). Right: multiple sequence alignment with A, B and C motifs and their corresponding logos. Blue and red regions indicate well-conserved residues.

RdRps existing in current databases but lacking a taxonomy (NA in Figure 4) were classified as Kitrino-, Duplorna- or Pisuviricota. As for the unknown RdRps (with bona-fide A, B and C motifs but absent from databases), one shared high sequence similarity with the Hypoviridae representative and was found in the same sample type (fungi from holm oaks), whereas the other was located in the vicinity of ssRNA(-) viruses (Chuviridae and Phenuiviridae) but shared no common origin with them.

## 4 Discussion

The study of environmental viromes has grown in popularity thanks to recent major advances in massive sequencing technologies. The scale of these analyses is highly variable, with some studies focusing on a particular species (Faizah et al., 2020; Wu et al., 2015), complete ecosystems (Hurwitz & Sullivan, 2013) or different environments (Daugrois et al., 2021; Ma et al., 2021). However, most of the advances in plant virology focus on crops or fruit trees of economic interest, which contrasts the scarcity of such studies in forestry (Rumbou et al., 2021). Reducing the scope even more to Mediterranean forests, with the Mediterranean basin being considered a biodiversity hotspot (Médail et al., 2019), hardly any information is found. In the present work, focused on 3 Mediterranean forest habitats in north-western Spain, 146 RdRps of viral origin have been found, of which 22 have no clear taxonomy yet and 2 are completely unknown. This shows the great lack of knowledge about viral diversity that exists in our own forests.

Most of the reported RdRps have been identified as ssRNA(+) viruses, which are accepted to be the evolutionary origin of the other two identified groups, ssRNA(-) and dsRNA (Koonin et al., 2015, 2021). Holm oak’s habitat showed the highest viral richness (57.5% of all the RdRps). Holm oak samples were taken in a “dehesa”, a unique ecosystem in the Iberian Peninsula that has been modified by man to simultaneously obtain livestock and non-timber forest resources. This makes dehesas an ecosystem with particularly high biodiversity (Moreno et al., 2016; Rodríguez-Rojo et al., 2022), so it is not surprising that there is also a high viral diversity that has never been investigated.

Some of the viral families have been found to be ubiquitous across sample types and habitats, which could be interpreted as a first indication of their cross-kingdom potential. This is the case for Botourmiaviridae, Partitiviridae, and Narnaviridae. Many genera in Botorumiaviridae family are known to infect fungi, and some of them have been identified on grapevine leaves affected by the oomycete *Plasmopara viticola* (Chiapello et al., 2020). Mycoviruses of this family can persist in their fungal host without the need for a capsid and are thought to require only the RdRp to replicate (Wang et al., 2020). Only the genus Ourmiavirus infects plants exclusively (Rastgou et al., 2009). The RdRp of this genus is most similar to that of the genera Mitovirus and Narnavirus (family Narnaviridae) and its movement protein (MP) is similar to that of the family Tombusviridae. The envelope protein bears some similarity to that of some plant or animal viruses. These viruses are very easily transmitted mechanically and no vector has been identified, so it is thought that there could be a horizontal transmission (Ayllón et al., 2020).

The Narnaviridae family has the simplest genome of all RNA viruses, encoding only one polypeptide in which RdRp is found. Within this family, the genus Narnavirus replicates in the cytoplasm, while Mitoviruses replicate in mitochondria and can cause hypovirulence in pathogenic fungi (de Rezende et al., 2021; Hillman & Cai, 2013). Phylogenetically, Narnaviruses are much more closely related to Ourmiaviruses (family Botourmiaviridae) than to Mitoviruses. Although no Mitoviruses have been found outside fungi, we’ve found one RdRp identified as a Mitovirus in a pine tree sample. Nonetheless, Mitovirus sequences have been found before in plant mitochondrial sequences (Hong et al., 1998; Marienfeld et al., 1999). As for the Partitiviridae family, many of its genera have characteristic hosts, which can be plants (Deltapartitivirus), fungi (Gammapartitivirus) and protozoa (Cryspovirus). Many sequences of this family have also been identified in arthropods (Cross et al., 2023). Here we’ve identified one genus, Betapartitivirus, whose hosts can be plants and fungi.

Regarding the family Tombusviridae, it has been described in animals (Yin et al., 2022), plants (Lappe et al., 2022) and fungi (Botella et al., 2022), matching our finding of this family in all sample types although restricted to holm oak habitat. Tombusviridae infections are usually limited to the root system, but they can infect the whole plant thanks to their MPs (Canuti et al., 2023). Tombusviridae are transmitted mechanically, by contact, through seeds or through a vector such as a fungus or an insect, depending on the genus. The Solemoviridae family infects plants, but it is known that one of its modes of transmission is also the use of insects as vectors (Sõmera et al., 2021), which may be one of the reasons why we have found this family only in arthropods.

For some families, only one RdRp has been detected in the samples analysed, but evidence of its ability to infect several kingdoms has been described. This is the case for the Totiviridae, Chuviridae, Phenuiviridae, Chrysoviridae, Polymicoviridae, Endornaviridae and two riboviruses with incomplete classification: the genus Botybirnavirus and the species Hubei picorna-like virus 15 (unclassified ribovirus). The family Totiviridae is associated with latent infections in fungi and protozoa. The families Chuviridae and Phenuiviridae, exclusive to insects, are the only ones that possess a ssRNA(-) genome. Many endogenous viral elements inserted within the host genome have been related to Chuviridae infections (Dezordi et al., 2020). The Chrysoviridae family infects fungi, plants and insects (Kotta-Loizou et al., 2020). The Polymycoviridae family affects fungi in which it causes hypovirulence or hypersensitivity to antifungals or bacteria, and is even capable of modulating carbon, nitrogen and iron metabolism (Kotta-Loizou & Coutts, 2022). Endornaviridae are ssRNA(+) viruses that do not have a true capsid as the genome is encapsulated together with a viral replicase and are able to infect plants and fungi. The genus Botybirnavirus (Orthornavirae) has fungi as natural hosts (Hough et al., 2023), however, we’ve found this RdRp in a holm oak tree sample, which could be indicative of some cross-kingdom capability.

Finally, some limitations of the present study should be noted. First, many sequences in viral databases have none or incomplete taxonomic assignment, and the high genetic variation of viruses makes it difficult to obtain an accurate assignment at genus or species level. Recurrent updating of the RdRps databases is necessary to facilitate the assignment of new ones. Secondly, although some of the virus families described may be “cross-kingdom” because they contain members that infect several kingdoms, it is more common for individual viruses to specialise in hosts of a single kingdom, as is the case, for example, with the Totiviridae family, where some genera infect fungi and others protozoa. Similarly, the *in silico* analysis performed does not allow us to verify whether finding RdRps in organisms from different kingdoms than expected implies that the host is susceptible to the virus, or that it only acts as a vector or asymptomatic carrier.

## 5 Conclusion

In this work we found great variation in the diversity of RNA viruses of declining Mediterranean forests, with holm oak forest having the highest richness and chestnut the lowest. Up to 15 viral families were identified overall, with Botourmiaviridae and Partitiviridae being the most abundant. In terms of genome type, ssRNA(+) viruses were the most represented, followed by dsRNA and two ssRNA(-) representatives. Viruses belonging to families with cross-kingdom capabilities such as Hypoviridae, Mitoviridae and Narnaviridae were also found. Lastly, two RdRP sequences had no matches in available viral databases and should be further investigated.

## 6 Data availability statement

Data and scripts used in this study are available in the following GitHub repository: https://github.com/serbiodh/2023_Virome_MedForests. Raw sequencing data are available at NCBI SRA under BioProject PRJNA1032577.

## 7 Author contributions

SDH, JNS and WAM collected the data. JNS and WAM carried out the laboratory procedures. SDH, PLPA and MP analysed the data. SDH, PLPA and JJD wrote the manuscript. SDH, JNS and JJD designed the sampling scheme. All authors contributed to the article and approved the submitted version.

## 9 Acknowledgements

Authors would like to thank Farooq Ahmad, Álvaro Benito Delgado, Irene Teresa Bocos Asenjo, Mariano Rodríguez Rey and Cristina Zamora Ballesteros for their valuable contributions to sampling.

